# Inference of Electrical Stimulation Sensitivity from Recorded Activity of Primate Retinal Ganglion Cells

**DOI:** 10.1101/2021.10.22.465478

**Authors:** Sasidhar S. Madugula, Ramandeep Vilkhu, Nishal P. Shah, Lauren E. Grosberg, Alexandra Kling, Alex R. Gogliettino, Huy Nguyen, Paweł Hottowy, Alexander Sher, Alan M. Litke, E.J. Chichilnisky

**Affiliations:** Department of Neurosurgery, Stanford University, Stanford, CA 94305, USA; Department of Electrical Engineering, Stanford University, Stanford, CA 94305, USA; Neurosciences PhD Program, Stanford University, Stanford, CA 94305, USA; School of Medicine, Stanford University, Stanford, CA 94305, USA; Santa Cruz Institute for Particle Physics, University of California, Santa Cruz, CA 95064, USA; Department of Ophthalmology, Stanford University, Stanford, CA 94305, USA; Hansen Experimental Physics Laboratory, Stanford University, Stanford, CA 94305, USA; Faculty of Physics and Applied Computer Science, AGH University of Science and Technology, Krakow, Poland; Facebook Reality Labs, Facebook, Mountain View, CA 94040, USA

## Abstract

High-fidelity electronic implants can in principle restore the function of neural circuits by precisely activating neurons via extracellular stimulation. However, direct characterization of the individual electrical responses of a large population of target neurons, in order to precisely control their activity, is often difficult or impossible. A potential solution is to leverage biophysical principles to infer sensitivity to electrical stimulation from features of spontaneous electrical activity, which can be recorded relatively easily. Here, this approach is developed and its potential value for vision restoration is tested quantitatively using large-scale high-density stimulation and recording from primate retinal ganglion cells (RGCs) *ex vivo*. Electrodes recording larger spikes from a given cell exhibited lower stimulation thresholds, with distinct trends for somas and axons, across cell types, retinas, and eccentricities. Thresholds for somatic stimulation increased with distance from the axon initial segment. The dependence of spike probability on injected current was inversely related to threshold, and was substantially steeper for axonal than somatic compartments, which could be identified by recorded electrical signatures. Dendritic stimulation was largely ineffective for eliciting spikes. These findings were quantitatively reproduced with biophysical simulations, and confirmed in tests on human RGCs. The inference of stimulation sensitivity from recorded electrical features was tested in simulated visual reconstruction, and revealed that the approach could significantly improve the function of future high-fidelity retinal implants.

## Introduction

Restoration of sensory capabilities using neural interfaces of the future will require evoking specific, naturalistic patterns of neural activity in large collections of neurons. This raises a crucial technical challenge: determining which electrodes in an electronic implant activate which cells and to what degree. In practice, with large neural populations, such measurements are difficult or even impractical. Thus, any properties of neurons that could be readily harnessed to reveal features of their electrical sensitivity could significantly enhance the capabilities of neural interfaces.

A potentially impactful approach would be to leverage knowledge of neuronal biophysics to *infer* how cells will respond to electrical stimulation, using only measurements of spontaneous electrical activity. Spontaneous activity can be recorded rapidly in parallel on many electrodes, and is not subject to electrical artifacts, and therefore can be analyzed relatively easily. For example, electrodes closer to a particular cell tend to record larger spikes, and correspondingly also require less current to evoke spikes. Thus, spike amplitudes recorded during spontaneous activity could potentially be used to infer electrical stimulation thresholds for every electrode and cell. This kind of inference has been suggested in theoretical and modeling studies, and experimental work in tissue culture (Boinagrov et al., 2010; Esler et al., 2018a; Fohlmeister et al., 1990; Loizos et al., 2014; Radivojevic et al., 2016; Tsai et al., 2012). However, no experimental tests of these ideas have been performed in functioning neural circuits. Thus, it remains unclear whether and how the electrical sensitivity of neurons in intact tissue can be reliably inferred and used effectively in a neural implant.

Here we propose an approach for electrical sensitivity inference in the context of a future high-resolution retinal implant, and empirically test its potential impact on vision restoration. We employed high-density multi-electrode recording and stimulation from hundreds of retinal ganglion cells (RGCs) in the primate retina *ex vivo* as a laboratory prototype, and probed the relationship between recorded spikes and sensitivity to electrical stimulation. Electrical sensitivity was systematically related to features of spiking activity, in a surprisingly consistent manner across hundreds of electrodes and many recordings, in both macaque and human retinas. This dependence exhibited systematically different properties based on the proximity of each electrode to the soma, axon, and dendrites of the cell, and on the cell type, and was in quantitative agreement with predictions from biophysical models. We show that these trends can be leveraged to accurately infer the responses of individual RGCs to electrical stimulation. Finally, we exploit these inferences to identify optimal electrical stimulation sequences for vision restoration, revealing the potential utility for a future high-fidelity retinal implant.

## Results

An experimental lab prototype of a future retinal implant was used to explore the relationship between recorded electrical features and electrical sensitivity of both macaque and human retinal ganglion cells (RGCs). Electrical recording and stimulation of RGCs in *ex vivo* isolated retinas was performed using a large-scale recording system (512 electrodes, 30 μm or 60 μm pitch, see Methods). After cell type identification by clustering responses to white-noise visual stimulation, 498 ON and OFF parasol and midget cells from 36 macaque retinal preparations, and 48 cells of the same types from 3 human retinas, were probed with electrical stimulation, eliciting individual, precisely-timed, directly-evoked spikes (Jepson et al., 2013; Sekirnjak et al., 2008). Next, to obtain a spatial signature of spontaneous spikes produced by each cell, the *electrical image* (EI) was computed and used to identify cellular subcompartments (see Methods). Finally, to determine the extracellular activation characteristics of each cell, responses evoked by electrical stimulation at each electrode were used to identify the *activation threshold* (current amplitude that evoked spikes with 50% probability), and the spatial map of the inverse of threshold over space was defined as the *electrical receptive field* (ERF) (Fig. 1).

**Figure 1.**
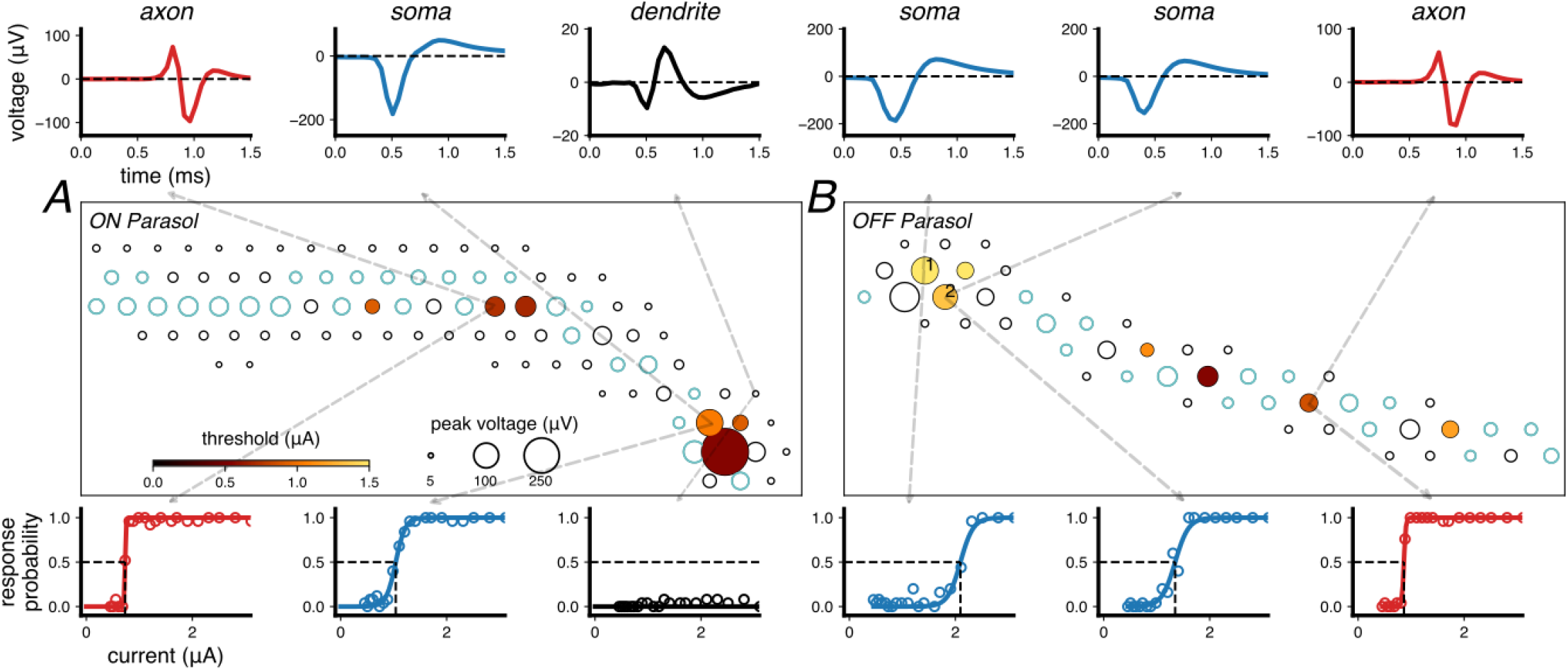
Electrical recording and stimulation of individual cells with a large-scale electrode array. A) ON parasol cell electrical image (EI) superimposed on the electrical receptive Field (ERF); circle area is proportional to recorded signal strength, and electrode color corresponds to electrical stimulation threshold. Electrodes that recorded spike amplitudes lower than the electrical noise threshold (30 μV) are not plotted, and those that did not evoke spikes over the current range tested are black and open. Electrodes for which the signal could not be analyzed due to axon bundle activity are light blue and open. Top: examples of recorded dendritic (black), somatic (blue), and axonal (red) spike waveforms (see Methods). Bottom: electrical activation probability as a function of current level at selected electrodes from cellular compartment (bottom). B) Similar to A, for an OFF parasol cell from the same retina.

### Features of electrical images correspond to the electrical receptive fields of ON and OFF parasol cells

Features of the EI, a relatively simple measurement, provided substantial information about the ERF, which summarizes electrical excitability over space but is far more difficult to obtain. Several observations about data from a single cell reveal trends that are probed in detail below. First, only electrodes that recorded large spike waveforms (Fig. 1, hollow circles) were able to evoke spiking (Fig. 1, filled circles) within the amplitude range tested (5-200 pC injected charge), although many such electrodes did not activate the cell at current levels below axon bundle activation threshold (Fig. 1, light-blue hollow circles; see Methods). Second, for the electrodes that did evoke a spike below bundle threshold (Fig. 1; see Methods), the EI amplitude at each electrode was related to the strength of the ERF: electrodes recording higher EI amplitudes (Fig. 1, circle size) also excited the cell more effectively (Fig. 1, circle color). The similarity between EIs and ERFs presumably reflects the fact that both spike amplitude and sensitivity to electrical stimulation are inversely related to the distance between the electrode and the cell (Rattay, 1987; Rattay et al., 2012; Sekirnjak et al., 2008). Third, somatic electrodes closer to the axon (Fig. 1B, electrode at circle 2) required less current to activate RGCs than somatic electrodes further from the axon (Fig. 1B, electrode at circle 1), even if the electrodes recorded spikes of similar amplitude (Fig. 1B, radii of circles 1 & 2). Fourth, electrical stimulation using dendritic electrodes typically did not elicit spiking within the tested stimulation amplitude range (Fig. 1A, top & bottom right). Finally, axonal electrodes exhibited steeper increases of firing probability in response to increasing stimulation current (Fig. 1 bottom, red vs. blue sigmoids, slopes), and recorded spike waveforms with smaller amplitudes (Fig. 1 top, red vs. blue waveforms, negative peak amplitudes), despite exhibiting activation thresholds in a range similar to those of somatic electrodes (0.2-2 μA). The latter suggested that the relationship between EIs and activation curves differs for somatic and axonal electrodes. In what follows, these trends in the relationship between the EI and ERF are explored across many cells and retinas.

### Electrical activation threshold is inversely related to recorded spike amplitude

The relationship between EI amplitude and ERF strength observed on axonal electrodes (Fig. 1, marker size and color) was consistent across cells. Recording and stimulation was tested on 61 electrodes overlying the axons of 53 ON and OFF peripheral parasol cells. Axonal thresholds exhibited a clear inverse relationship to EI amplitudes (Fig. 2 A&B, red markers). To determine whether this relationship holds across preparations, recording and stimulation features from 152 axonal electrodes were compared across recordings (Fig. 2C). A consistent inverse relationship was observed for electrodes overlying axons (Fig. 2C, circle and square markers respectively), similar to the trend observed in a single retina (Fig. 2 A&B, red points vs. points in C). The relationship between axonal EI amplitudes and activation thresholds was fitted by an inverse function based on studies indicating that electrode distance is linearly related to activation threshold for electrodes less than 50 μm away (Rattay and Wenger, 2010): 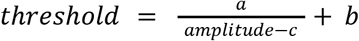 (*equation 1*, Fig 2C, dashed line, see Methods).

**Figure 2.**
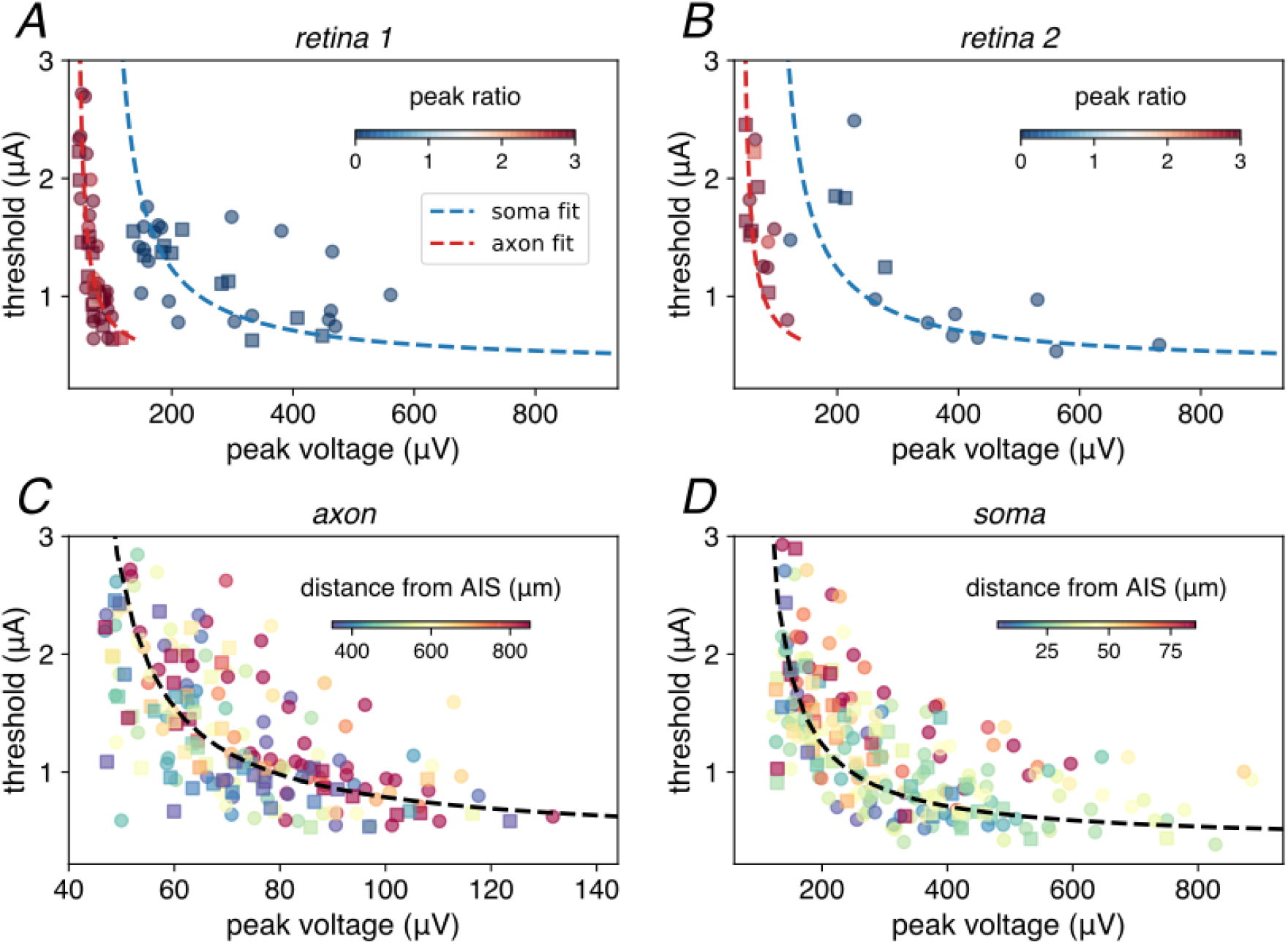
Relationship between activation thresholds, electrode location, and EI amplitudes for peripheral macaque parasol cells. In all plots, points corresponding to ON Parasol cells are denoted by circular markers, and OFF Parasol cells by square markers. A,B) Electrical activation threshold vs. recorded spike amplitude for axonal (red) and somatic (blue) electrodes in two different retinal preparations (see Methods). Colored dashed lines indicate maximum likelihood curve fits to the aggregated data for somas (blue, fit to data in C) and axons (red, fit to data in D, parameters: axon, equation 1 - a = 6.5, b = 0.4 μA, and c = 35 μV, soma, equation 2 - a = 22, b = 0.4 μA, c = 96 μV, d = 0.0146, and e = −0.428, see Results). Aggregated thresholds vs. EI amplitudes for axonal (C) and somatic (D) electrodes, collected from 23 retinas; dashed lines correspond to curve fits to the data. Color indicates distance from the axon initial segment (AIS) location.

Similarly to axonal activation thresholds, somatic thresholds exhibited an inverse dependency on EI amplitude, as observed in 55 somatic electrodes overlying 53 ON and OFF parasol cells in two retinal preparations (Fig. 2 A&B, blue markers). However, pooling the recording and stimulation features and inspecting the locations of 94 somatic electrodes revealed that somatic activation thresholds depended on the distance of the stimulating electrode from the putative location of the axon initial segment (AIS, Fig. 2D, marker colors; see Methods). Specifically, for a given RGC, somatic electrodes near the AIS (Fig. 2D, blue markers below dashed line) required less current to cause activation than electrodes farther away (Fig. 2D, red markers above dashed line), despite recording spikes with similar peak amplitudes. Quantitative characterization of this trend revealed that the fractional difference between measured somatic activation thresholds and estimates based solely on EI amplitude increased roughly linearly as a function of electrode distance from the AIS (r^2^ = 0.78). Taken together, these findings suggested the following equation for estimating somatic activation thresholds from recorded somatic spikes: 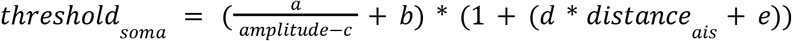 *(equation 2)*. Note that axonal electrodes did not exhibit any discernible dependence on distance to the AIS (Fig. 2C, marker colors) or to the midline of the axon (see Methods).

Across many cells, electrodes near dendrites did not reliably elicit spikes (Fig 1A bottom, black trace). On average, only 9% of identified electrodes recording significant spikes (see Methods) from each cell were identified as recording from dendrites (3577 cells, 34 retinas). Of these electrodes, none were able to elicit spikes with probability greater than 0.5 over the range of amplitudes tested, compared to 74% for all axonal electrodes and 93% for all somatic electrodes (see Methods). Thus, the dendritic data were insufficient to explore the EI amplitude-activation threshold relationship and were not further analyzed.

### Activation curve slope is inversely related to activation threshold

A complete characterization of RGC electrical responsivity entails determining not only the 50% activation threshold, but also the spiking probability over a range of stimulus amplitudes--i.e. a full *activation curve* (Fig. 3A). A simple possibility is that, for each cellular compartment, the activation curve has a stereotyped form across cells, because of the similarities in morphology and membrane dynamics between cells, and that this form scales with the injected current according to the electrical impedance between the electrode and the site of activation. In this case, normalizing measured activation curves by their associated threshold should result in a constant form. Indeed, normalization revealed strikingly similar activation curves across retinal preparations for all somatic electrodes, and for all axonal electrodes, but very different forms for the two compartments (Fig 3C, blue vs. red curves): axonal activation curves featured a relatively steep “all-or-none” relationship, while somatic activation curves were shallower.

**Figure 3.**
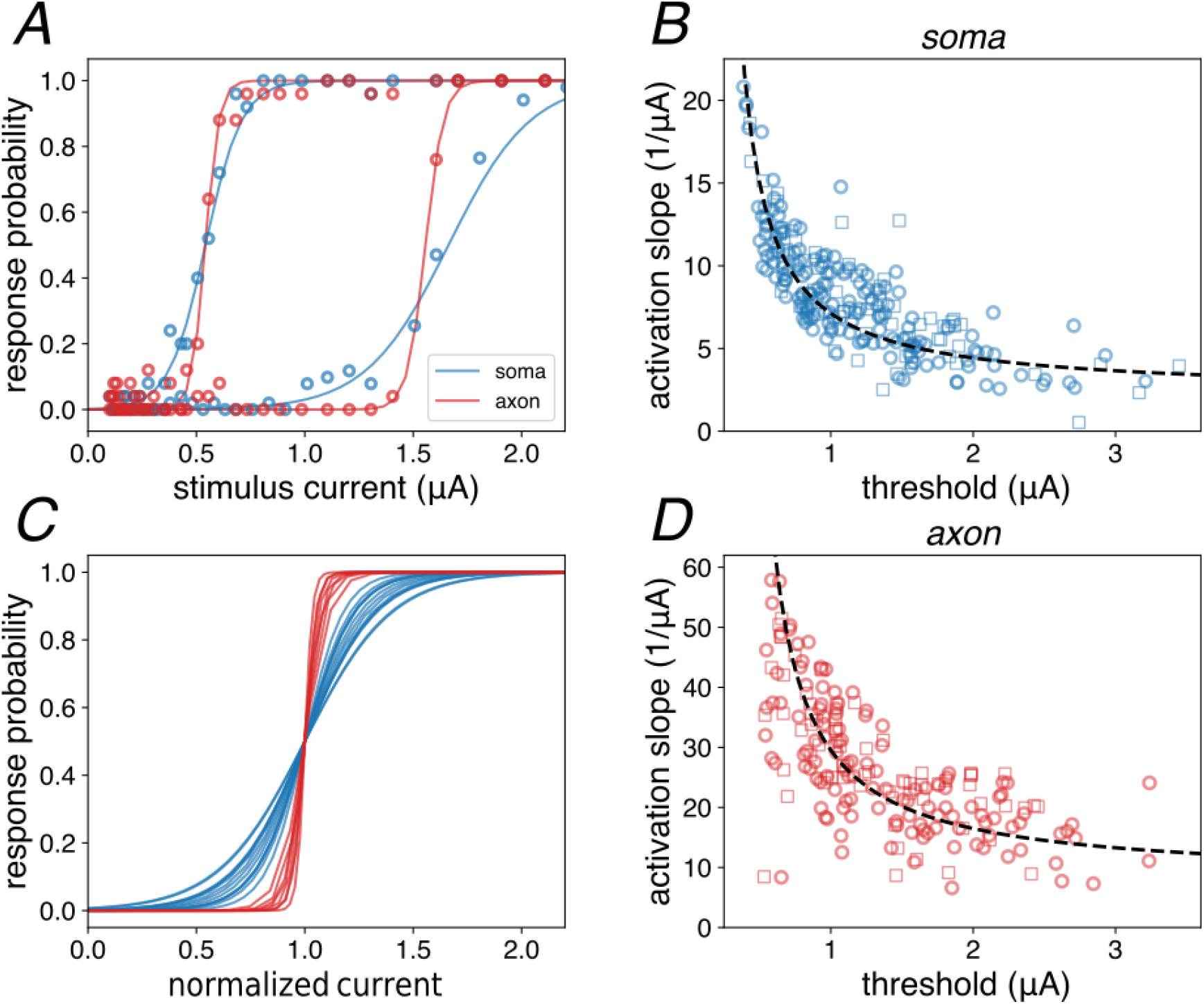
Relationship between activation curve slopes and thresholds. A) Two somatic (blue) and two axonal (red) measured sigmoidal activation curves. C) Activation curves shown in (A) along with 10 additional examples for each compartment, after current values were normalized by each curve’s activation threshold (see Results). Activation curve slopes vs. thresholds for somatic (B) and axonal (D) electrodes, with curve fits (dashed line), with parameters = 3.94 for somas and 4.91 for axons, b = 2.25 for somas and 8.72 for axons, and c = 0.19 for somas and 0.6 for axons (see Results). Fitted activation curves with spuriously high slopes resulting from poorly estimated spiking probabilities were excluded (see Methods).

The similarity of normalized activation curves within each cellular compartment indicates that the slope is inversely related to the threshold. To characterize this relationship further, activation curve slopes were examined as a function of activation threshold, separately for 94 somatic and 152 axonal electrodes (Fig. 3 B&D). This relationship was fitted by an inverse function: 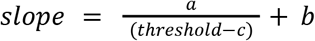 (*equation 3*, Fig. 3 B&D, dashed lines, for fit parameters see caption). For each compartment, ON and OFF parasol cells exhibited a similar relationship between activation curve slope and threshold (Fig. 3 B&D, circular and square markers, respectively). The high fidelity of these relations (r^2^ = 0.82 for somas and 0.74 for axons) suggests that characterizing the ERF of a cell (which only reflects threshold at each electrode) is sufficient to reconstruct the full set of activation curves for the cell at each electrode.

### Electrical recording and activation properties generalize across cell types, retinal location, and species

The relationships observed between EI amplitude, electrode position, and activation threshold for peripheral macaque parasol cells originate from their biophysical properties, which could potentially generalize to other RGC types, to RGCs in the central retina, and to human RGCs. To test this hypothesis, electrical recording and stimulation of the somas and axons of peripheral midget cells, parasol cells in the central *raphe* region, and peripheral midget and parasol cells from human retinas were examined (see Methods, Fig. 4).

**Figure 4.**
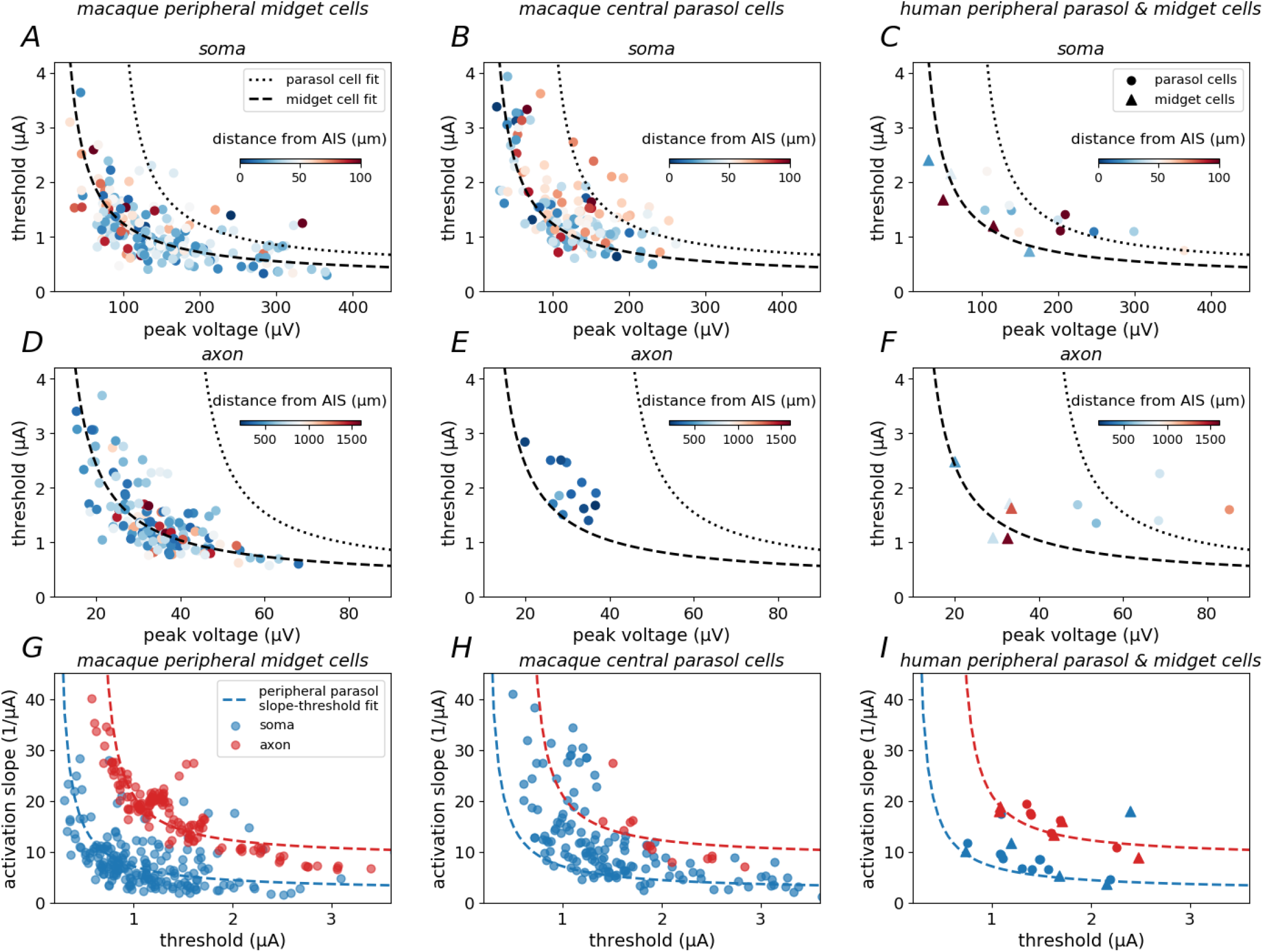
Relationship between activation thresholds, activation slopes, electrode location, and recorded spike amplitudes for RGCs of different types, eccentricities, and species. In panels A-F, marker color corresponds to distance from estimated axon initial segment (AIS) location; dashed lines indicate fits to parasol cell data from Fig. 2, dotted lines correspond to fits to midget cell data in panels A&D (parameters: a = 94.2, b = 0.23 μA, and c = 7 μV for somas, and a = 23.4, b = 0.28 μA, and c = 9 μV for axons, equation 1). A,D) Thresholds vs. spike amplitudes for peripheral midget cell somatic (top) and axonal (bottom) electrodes aggregated from 15 macaque retina recordings(circular markers). B,E) Thresholds vs. spike amplitudes for macaque central parasol somatic (top) and axonal (bottom) electrodes collected from 10 different macaque retina recordings. C,F) Thresholds vs. spike amplitudes for peripheral human parasol (circles) and midget (triangles) cell somatic (top) and axonal (bottom) electrodes from 3 human retina recordings. G-I) Relationship between activation curve slopes and activation thresholds for macaque peripheral midget (G), macaque central parasol (H), and human peripheral parasol (circles) and midget (triangles) cells (I). Dashed lines correspond to the somatic (blue) and axonal (red) slope-threshold relationships in Fig. 3 B,D. Blue denotes somatic electrodes, and red denotes axonal electrodes.

Peripheral macaque midget cells exhibited a clear inverse relation between recorded EI amplitudes and activation thresholds (Fig. 4 A&D), for both somas and axons, which were characterized by a translated version of the relation obtained for peripheral parasol RGCs (equation 1; Fig. 4 A&D, dashed vs. dotted line, r^2^=0.83 for somas, 0.79 for axons). However, midget RGC somatic electrode thresholds were not obviously influenced by proximity to the AIS in human or macaque (r^2^ = 0.0001, Fig. 4B, marker colors, see Discussion). Central parasol cell activation thresholds, measured using a MEA with 30 μm pitch (instead of 60 μm pitch used for all other experiments), also depended on recorded EI amplitudes, with inverse relationships that were well-described by the fit to peripheral midget RGCs (Fig. 4 B&E, markers vs. dashed curve, r^2^=0.76 for somas, 0.52 for axons). The similarity of electrical recording and activation features between central parasol cells and peripheral midget cells may arise from the similarity in cell size (Watanabe and Rodieck, 1989). In contrast, however, the activation thresholds of somatic electrodes overlying central parasol cells depended on proximity to the AIS (r^2^ =0.64, linear fit slope: 0.005, intercept: 0.03, Fig. 4B marker colors), echoing the trend seen in peripheral parasol cells (Fig. 2, fit parameters *d* and *e*).

Finally, to explore the possibility of applying the above findings in a clinical implant, the EI amplitudes and activation thresholds of human peripheral parasol and midget cells were examined. While only a few cells were analyzed due to limited tissue availability and avoiding axon bundle activation (see Methods), EI amplitudes and activation thresholds for somatic and axonal electrodes broadly coincided with the curve fit obtained using peripheral macaque parasol cells (equation 1, Fig. 4 A&D, markers vs. dashed curve).

The dependence of activation curve slope on activation threshold may also generalize to different RGC types, eccentricities, and to the human retina in a similar manner. In fact, the slope-threshold relationships for macaque peripheral midget cells, macaque central parasol cells, and human peripheral parasol and midget cells were well-characterized by the inverse activation curve slope-threshold relationships observed for the somatic and axonal electrodes for macaque peripheral parasol cells (Fig. 3), with mean r^2^ values of 0.78, 0.68, and 0.61 respectively (Fig. 4 G,H,I blue and red dashed lines vs. corresponding markers).

### Biophysical simulations corroborate experimentally observed extracellular activation trends

Simulated extracellular stimulation of mammalian RGCs (Fohlmeister et al., 2010; Fohlmeister and Miller, 1997) reproduced the main empirically observed trends in electrical activation properties. Simulations were conducted using the NEURON software package (Carnevale and Hines, 2006, see Methods) to implement a previously published electrical circuit model of a rat RGC, with soma and axon diameters modified to reflect parasol and midget cell geometry, based on morphological measurements from primate RGCs (see Methods, Fig. 5A. The model cells were simulated in a uniform medium with isotropic resistivity. To simulate nondeterministic activation, spontaneous membrane voltage fluctuations (mimicking noise channel noise and synaptic inputs) were modeled as Gaussian noise in each model segment (see Methods). Stimulating electrodes were modeled as a row of point current sources with 60 μm spacing along the cell and axon, at a retinal depth 7 μm from that of the axon (Fig. 5A, black, blue, and red row of bars spanning the top). Voltages at each simulated electrode were obtained by summing the voltage fluctuations resulting from action potentials in the model segments <u>(Gold et al. 2006)</u>.

**Figure 5.**
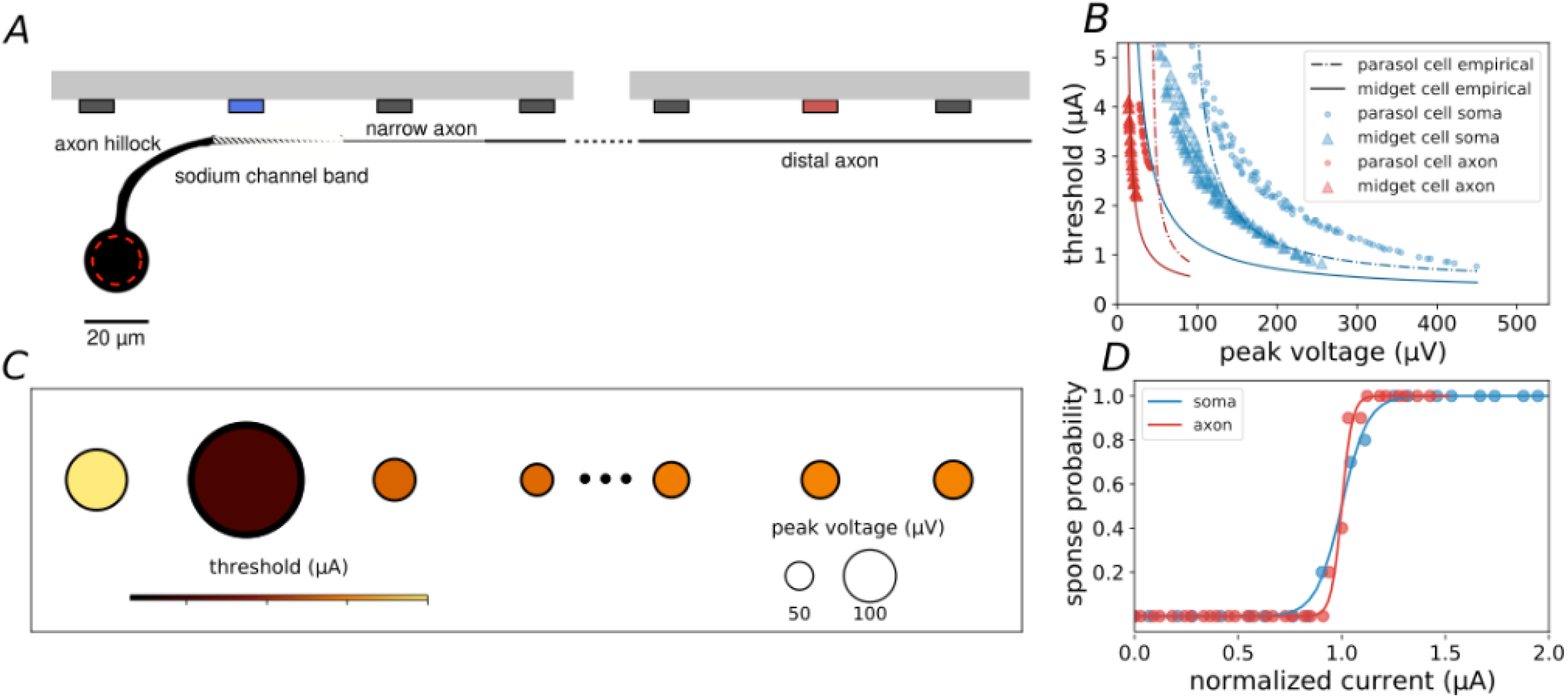
Simulated extracellular activation properties of RGCs. A) Finite-element NEURON model schematic with parasol cell soma (red dashed outline corresponds to midget cell soma, note that thickness of all other compartments also differs for midget cells but is not shown, see Methods), axon hillock, sodium channel band, narrow axon, and distal axon compartments, along with rectangles representing overlying simulated electrode positions, including the somatic (blue) and axonal (red) electrodes used in B&D. Ellipsis indicates spatial discontinuity to allow for visualization of the distal axon. B) Activation thresholds vs. recorded spike amplitudes for simulated (circular and triangular markers) and experimentally collected (dash-dotted and solid lines) data, for somatic (blue) and axonal (red) electrodes overlying parasol (circles) and midget (triangles) cells. C) Simulated parasol cell ERF with electrode locations corresponding to each electrode x-position in (A). Circle sizes represent peak recorded spike amplitudes, colors indicate activation thresholds. D) Simulated somatic (blue) and axonal (red) activation curves.

Analysis of these simulations matched several of the key findings in the recorded data. First, simulated electrodes located over the soma (Fig. 5A, blue rectangle) recorded spikes with 3-5 times the amplitude of electrodes overlying the axon (Fig. 5A, blue rectangle), but activated the model cell with similar current thresholds (Fig. 5B, blue vs. red markers). Second, stimulation and recording over a range of electrode-cell distances (see Methods) resulted in inverse relationships between spike amplitude and activation threshold for both somatic and axonal electrodes, exhibiting significant overlap with the corresponding experimental data for both parasol and midget cells (Fig. 5B, red and blue circular and triangular markers vs. dashed and solid lines, see Discussion). Third, electrodes recording somatic spike waveforms closer to the AIS had lower activation thresholds than electrodes farther from the AIS (Fig. 5C, circle colors). Finally, stimulation with increasing current produced steeper activation curves at axonal electrodes than at somatic electrodes (Fig. 5D, blue and red curves). Taken together, these results suggest that the experimentally observed relationships between the electrical features of RGCs and their activation properties are mediated by well-understood biophysical mechanisms.

### Activation curves can be accurately inferred from electrical images

The systematic relationship between the EI and ERF probed above proved useful for inferring the parameters of ON and OFF parasol and midget cell activation curves using only recorded spikes. To test this, the previously fitted inverse relations were used to estimate thresholds, which were then compared to measured thresholds, for the somas and axons of 102 parasol cells (Fig. 6A, example ON Parasol cell), across 246 electrodes from 23 different preparations (Fig. 6 B,C,E,F). The accuracy of inferred thresholds was compared to that of a single, naive estimate of the activation threshold obtained by averaging measured activation curve parameters over axonal and somatic electrodes across 53 retinal preparations (Fig. 6 B,C,E,F horizontal dashed lines, average threshold 1.2 μA, average slope 12.1 1/μA). Inferred ERFs were qualitatively similar to measured ERFs for individual cells (Fig. 6A left vs. right, circle colors). Across axonal electrodes, 56% of activation thresholds were estimated within 25% of the measured value, compared to 33% obtained by averaging (Fig. 6E). Across somatic electrodes, 48% of inferred thresholds were within 25% of the measured value, compared to 28% obtained by averaging (Fig. 6B). Activation slopes inferred from these estimated activation thresholds were within 25% of the measured slope value for 58% of axonal electrodes, compared to 14% using the average slope (Fig. 6F), and for 67% of somatic electrodes, compared to 22% using the average slope (Fig. 6C).

**Figure 6.**
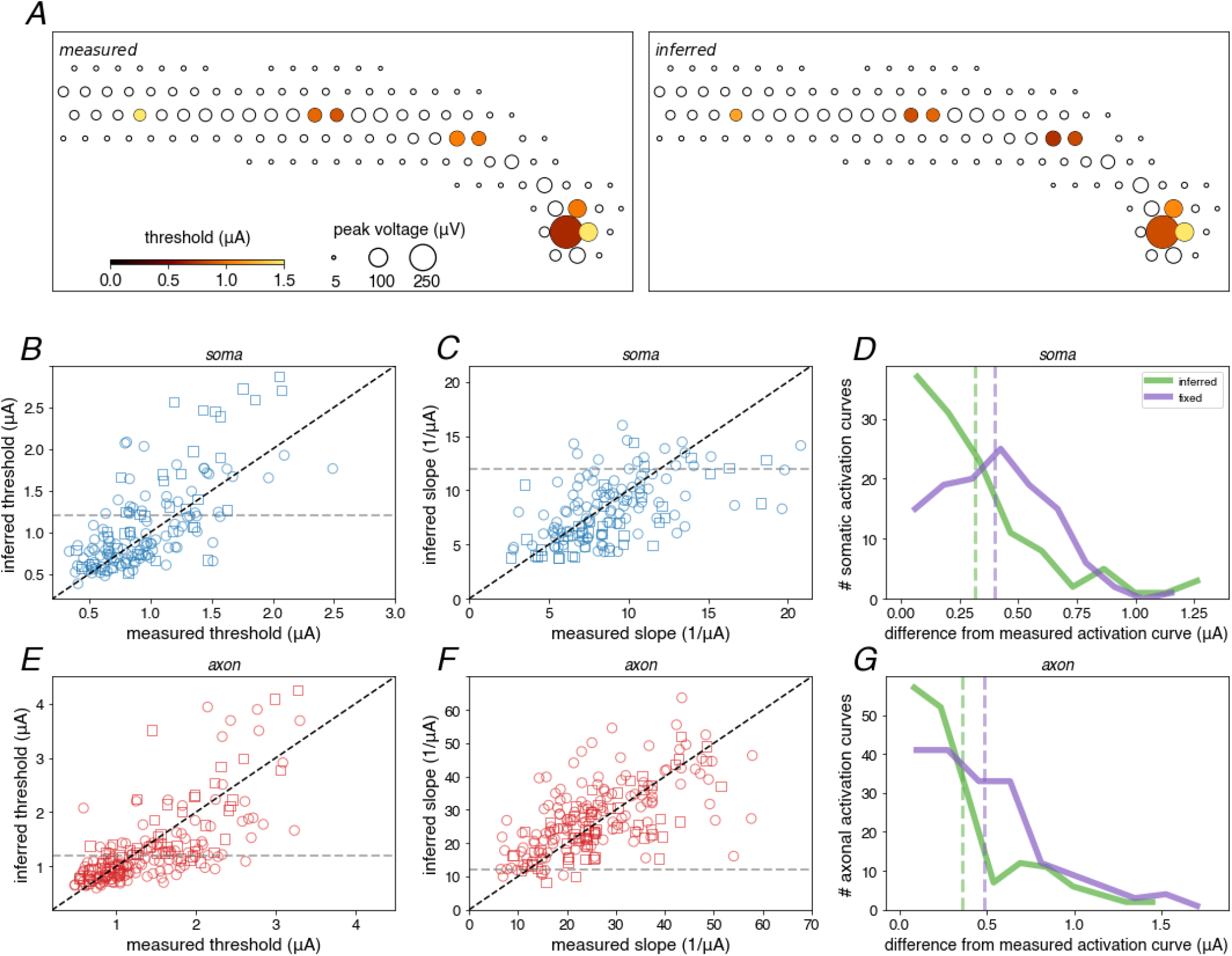
Comparison of inferred and measured activation properties. A) Measured vs. inferred ERFs for a single ON parasol cell. Circle size corresponds to spike amplitude recorded at each electrode; circle shade corresponds to measured (left) or inferred (right) activation threshold at each electrode. Inferred vs. measured somatic (B) and axonal (E) thresholds, and slopes (C,F), pooled across 259 cell-electrode pairs. ON and OFF parasol cells are indicated by circular and square markers, respectively. Dashed line corresponds to x=y. D) Histograms of the absolute differences between integrals of inferred (green), or fixed (purple) somatic activation curves, with respective mean values indicated by vertical dashed lines. G) Histograms of absolute differences between integrals of inferred and measured axonal activation curves.

These inferred activation curves provided accurate estimates of cellular activation probabilities over the range of applied stimulus currents. This was determined by pooling data across 246 electrodes and comparing measured curves to the corresponding inferred curves (using the fitted inverse relations for threshold and slope inference, respectively), or to a single, naively estimated sigmoidal activation curve derived from average parameters (same as above). The difference between each measured and estimated sigmoidal activation curve was estimated by computing the absolute difference of the integrated area between the curves. The separation between fitted and inferred activation curves was significantly smaller than the separation between fitted and averaged curves for somas and axons (Fig. 6 D&G, Wilcoxon signed rank test between green and purple histograms: p=0.002 for somas, p=0.02 for axons). The mean activation curve separation for inferred somatic activation curves was 22% closer to zero than for the fixed curve, and 30% closer for axons (Fig. 6 D&G, vertical dashed lines).

### Inferred activation curves can be used to inform stimulation choices for vision restoration

The inferred electrical responsivity of RGCs (Fig. 6) was useful for optimizing electrical stimulation over space using an algorithm designed for use in a high-resolution epiretinal implant (N. P. Shah et al., 2019). This was demonstrated by estimating the cumulative visual perception (or *image reconstruction*, (Brackbill et al., 2020; Warland et al., 1997)) resulting from a collection of current pulses delivered by electrodes across the array (see Methods), separately in three different retinal preparations. In each preparation, activation of each ON or OFF parasol cell was assumed to contribute a particular image component to visual perception, corresponding to the optimal linear reconstruction filter obtained using responses to white-noise visual stimuli (see Methods). Spiking probabilities were computed using either measured responses to electrical stimulation (ground truth), inferred responses based purely on electrical recordings, or a fixed activation curve representing the average across 246 electrodes and 102 cells in 53 preparations. Each image was translated into a spatial pattern of stimulation, using 40 individual electrodes and current amplitudes selected to minimize the normalized, pixel-wise mean-squared-error (NMSE, see Methods) between the reconstructed and true image.

Qualitative comparison between reconstructed images revealed that choosing electrodes based on inferred activation curves resulted in substantially more accurate reconstruction than could be obtained by using the fixed curve, and approached the best achievable reconstruction obtained using measured activation curves (Fig. 7A, example reconstructions of two target images in a single preparation). Comparing the accuracies of 45 reconstructed targets across 3 retinal preparations revealed that choosing stimuli using inferred activation curves provided a 51% improvement in NMSE over using fixed curves (see Methods, Fig. 7B, dashed red, green, and black dashed line slopes). These results suggest that inference of activation curves from EIs has the potential to produce more accurate visual perception with a future, closed-loop epiretinal implant.

**Figure 7.**
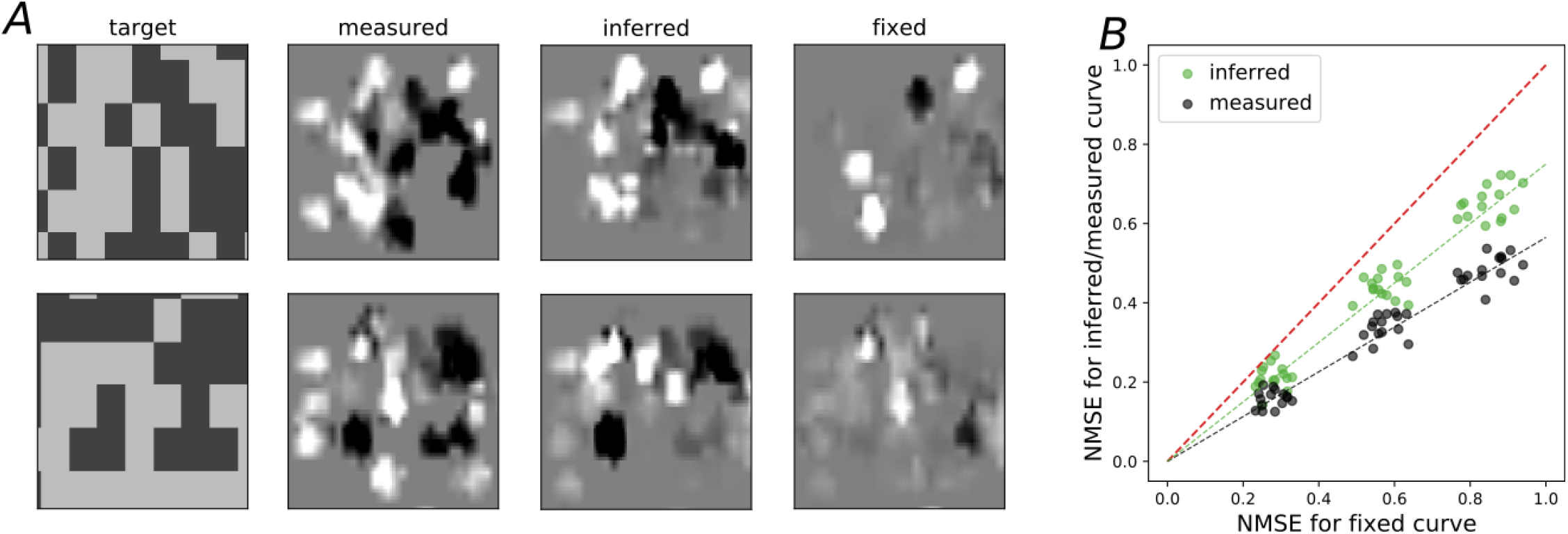
Simulated image reconstruction using inferred, measured, and fixed activation curves. A) Reconstructed images in one retinal recording for two targets (top and bottom rows respectively, left), using measured (middle-left), inferred (middle), and fixed control (middle-right), activation curves. B) Normalized mean-squared-error (NMSE) image reconstruction for inferred vs. fixed (green markers) and measured vs. fixed (black markers) across 15 targets and 3 retinal recordings Data from each recording forms a cluster of points. Red dashed line indicates x=y, green dashed line represents linear fit to inferred vs. fixed points, black dashed line represents linear fit to measured vs. fixed points.

## Discussion

The present data reveal a systematic relationship between the electrical features of neurons and their responsivity to electrical stimulation, most notably an inverse relationship between spike amplitude and stimulation threshold for each cellular compartment. These trends were replicated in simulations with a biophysical model, indicating that they arise from well-understood properties of neurons. Finally, we showed that the observed relationships can be harnessed to accurately infer the extracellular activation properties of cells over space, and thus may be useful in guiding electrical stimulation choices relevant to prosthetic vision.

Several key features of electrical activation properties presented here are supported by previous studies. First, the inverse relationship between spike amplitude and activation threshold agrees with theoretical predictions. These predictions are based on the assumption that extracellular activation depends on the second spatial derivative of the voltage deflection along the membrane of the axon (Rattay, 1987; Rattay and Wenger, 2010), and suggest that the distance-threshold relationship is linear at electrode distances less than ∼50 μm. The linear relationship in turn supports the observed inverse relationship between recorded spike amplitude and threshold. In addition, reports from previous experimental and theoretical studies indicate that the AIS, which is positioned between the soma and distal axon of RGCs, is highly excitable (Boiko et al., 2003; Esler et al., 2018b; Fried et al., 2009; Radivojevic et al., 2016; Rattay and Wenger, 2010; Werginz et al., 2020), consistent with the observed increase in activation threshold with increasing distance from the AIS for electrodes near the soma. However, applying the inference relations presented here to other devices may require the collection of at least some electrical stimulation data, despite the putative extension of the biophysical trends to all standard neuron types, because stimulation thresholds (but not recorded spike amplitudes) may vary based on electrode geometry (Cao et al., 2015; Lempka et al., 2011; Viswam et al., 2019).

In a notable previous study, inference of activation thresholds from the recorded features of electrically evoked spikes was performed in 14 cultured cortical neurons (Radivojevic et al., 2016). Inference was performed using multivariate regression on eight features of recorded spikes, revealing that the two critical features for inference were spike amplitude and time of spike onset. These two features roughly correspond to those used for threshold inference in the present work: spike onset times often identify the cellular compartment, due to the stereotyped time course of action potential propagation.

Several limitations of the present work bear mentioning. First, the results were obtained from healthy primate retinas, but degenerate retinas would be more relevant for application to epiretinal implants. Experimental and computational studies of electrical stimulation using rat models of retinal degeneration indicate that recorded spike waveforms and stimulation thresholds do not significantly change with degeneration (Loizos et al., 2018; Sekirnjak et al., 2009). However, other studies employing stimulation using long pulses and large electrodes in mouse models suggest that thresholds increase (Jensen and Rizzo, 2008; O’Hearn et al., 2006; Suzuki et al., 2004). It remains to be seen whether the relationship between recorded spikes and activation thresholds is the same in diseased and healthy primate retina.

Second, for each electrode, only activation thresholds below axon bundle threshold were included in this study. Estimating axon bundle thresholds currently requires electrical stimulation and collection of electrical response data. In theory however, axon bundles could be removed during device implantation on the retina by incising the tissue on the distal edge, inducing Wallerian degeneration.

Third, somatic activation thresholds for peripheral midget cells did not exhibit the inverse dependency on proximity to AIS observed for peripheral and central parasol cells. This could be due to the difficulty of accurately estimating the location of the AIS in small cells with an MEA featuring 60 μm electrode pitch, since the trend is clear in central parasol cells, which have EIs that are roughly the same size as peripheral midget cells. Conversely, it may be the case that there is a true difference in AIS sensitivity of parasol and midget cells, though this observation is not suggested by any previous findings.

Finally, although the biophysical model (Fohlmeister et al., 2010; Fohlmeister and Miller, 1997) captured general trends in the data, it did not exactly quantitatively reproduce the experimental results (Fig. 5). This discrepancy may reflect several factors. The properties of the model cell are derived from rat ON-alpha RGCs, which may differ from primate parasol RGCs in crucial features such as cell size and channel density. The model also does not distinguish between the various subtypes of sodium channels in distinct cellular compartments (Boiko et al., 2003; Van Wart and Matthews, 2006). Additionally, the resistivities of the retinal layers in primates is unknown, and reported measurements vary significantly (Abramian et al., 2015; Kasi et al., 2019; Wang and Weiland, 2015). Lastly, modeling the non-deterministic nature of extracellular activation, which presumably results from membrane voltage fluctuations caused by ion channels and synaptic inputs, relies on patch clamp data that can be obtained reliably from RGC somas but not from axons. For these reasons, the model was used only to generally confirm the relationships observed in the empirical data, and suggest possible biophysical mechanisms. For example, one potentially interesting insight derived from the model is that the expected difference in the recorded spike amplitude - activation threshold relationship between parasol and midget cells was produced simply by changing the overall cell size and not the sodium channel distribution.

Despite these limitations, inference of RGC activation properties purely from spontaneous activity using biophysically-grounded relationships has the potential to improve the function of a future retinal implant, by wholly or partially eliminating the need to perform painstaking calibration of a large-scale retinal interface. This is significant because electrical stimulus calibration generally requires sequential testing of each individual electrode, which is time-consuming, and more importantly is greatly complicated by electrical artifacts, which make it difficult to identify spikes evoked by electrical stimulation. These requirements could pose substantial technical challenges in the clinical setting, and highlight the value of the inference approach described. A hybrid approach could entail performing full electrical calibration only for cells with small recorded spikes, to supplement the more reliable inferred electrical activation curves of cells with large recorded spikes. Alternatively, initial estimates of electrical thresholds obtained from recording alone, or using inferred sensitivity as priors in Bayesian estimation (N. Shah et al., 2019), could be used to more quickly identify thresholds using limited electrical stimulation and recording. Inference also allows for rapid re-calibration of electrical stimulation over time, potentially compensating for small movements of an implanted device. Furthermore, the similarity between the electrical impedance and neuronal activation characteristics of the retina and other neural circuits (Branner et al., 2001; Hillman et al., 2003; Rattay, 1987; Royeck et al., 2008), and the surprising degree to which broad biophysical principles consistently apply to different populations of RGCs in intact tissue, suggests that the ability to infer electrical activation properties from extracellular recordings may be possible in other neural circuits and thus could be generally relevant for neural implants.

## Acknowledgements

Human eyes were provided by Donor Network West (San Ramon, CA). We are thankful for the cooperation of Donor Network West and all of the organ and tissue donors and their families, for giving the gift of life and the gift of knowledge, by their generous donations. We thank J. Carmena, T. Moore, W. Newsome, M. Taffe, S. Morairty, J. Horton, and the California National Primate Research Center for providing macaque retinas. We thank K. Jensen, M. Allen, and K. Tinajero, K. Berry, K. Williams, B. Morsey, J. Frohlich, and M. Kitano for helping obtain access to macaque retinas and metadata. We thank D. Palanker, F. Rieke, S. Mitra, P. Li, T. Carnevale, M. Hines, F. Rattay, P. Tandon, E. Wu, R. Vilkhu, V. Fan, M. Zaidi, C. Rhoades, N. Brackbill, G. Goetz, S. Wienbar, and the Stanford Artificial Retina team for helpful discussions. We thank K. Kish and J. Weiland for helpful discussions and graciously sharing the base code for the biophysical modeling simulations used in this work. We thank H. Nguyen for contributions to electrical spike sorting in the central retina. We thank R. Samarakoon and S. Kachiguine for technical assistance. We thank L. Jepson, P. Li, M. Greshner, G. Field, J. Gautier, A. Heitman, and G. Goetz for participating in data collection. Support: NIH NEI F30-EY030776-03 (SM), Polish National Science Centre grant DEC-2013/10/M/NZ4/00268 (PH), Pew Charitable Trust Scholarship in the Biomedical Sciences (AS), a donation from John Chen (AML), Research to Prevent Blindness Stein Innovation Award, Wu Tsai Neurosciences Institute Big Ideas, NIH NEI R01-EY021271, NIH NEI R01-EY029247, and NIH NEI P30-EY019005 (EJC).

## Author Contributions

Conceptualization, S.S.M., L.E.G., and E.J.C.; Methodology, S.S.M., L.E.G., N.P.S., and E.J.C.; Investigation, S.S.M., L.E.G., N.P.S., A.K., and A.R.G.; Writing – Original Draft, S.S.M. and E.J.C.; Writing – Review & Editing, S.S.M. and E.J.C.; Funding Acquisition, S.S.M., P.H., A.S., A.M.L., and E.J.C.; Resources, S.S.M., L.E.G., N.P.S., A.K., A.R.G., P.H., A.S., A.M.L., and E.J.C.; Supervision, E.J.C.

## Funding

NIH NEI F30-EY030776-03 (SM)

Polish National Science Centre grant DEC-2013/10/M/NZ4/00268 (PH)

Pew Charitable Trust Scholarship in the Biomedical Sciences (AS)

a donation from John Chen (AML)

Stanford Medicine Discovery Innovation Award (EJC)

Research to Prevent Blindness Stein Innovation Award (EJC)

Wu Tsai Neurosciences Institute Big Ideas (EJC)

NIH NEI R01-EY021271 (EJC)

NIH NEI R01-EY029247 (EJC)

NIH NEI P30-EY019005 (EJC)

## Methods

### Experimental setup

Custom 512-electrode and 519-electrode systems (Grosberg et al., 2017; Hottowy et al., 2012, 2008) were used to stimulate and record peripheral and central retinal ganglion cells (RGCs), respectively, in isolated rhesus macaque monkey retinas (*Macaca mulatta*) and human retinas. Human eyes were obtained from three brain-dead donors (29 year-old hispanic male, 27 year-old hispanic male, 47 year-old caucasian female) through Donor Network West (San Ramon, CA). Macaque retinas were obtained from terminally anesthetized animals euthanized during the course of research performed by other laboratories, in accordance with institutional and national guidelines and regulations. Ocular hemisection was performed in ambient indoor lighting following enucleation, the vitreous was removed, and the posterior portion of the eye was stored in warm, oxygenated, bicarbonate buffered Ames’ solution (Sigma-Aldrich) in darkness. Patches of retina ∼3 mm on a side were isolated under infrared light, placed RGC side down on the multielectrode array, and superfused with Ames solution at 35°C. Electrodes were 8-15 μm in diameter and were electroplated with platinum. The multi-electrode arrays (MEAs) consisted either of 512 electrodes arranged in a 16 × 32 isosceles triangular lattice with 60 μm spacing, or of 519 electrodes arranged in a 16 × 32 isosceles triangular lattice with 30 μm spacing (Litke et al., 2004). Voltage recordings were band-pass filtered between 43 and 5,000 Hz and sampled at 20 kHz. Spikes in the voltage recordings from individual RGCs evoked by light stimulation (which produced no electrical artifacts) were identified and sorted using previously described spike sorting techniques (Litke et al., 2004).

### Visual stimulation and cell type classification

To identify the type of each recorded cell, as well as the location and shape of the visual receptive field (RF), the retina was visually stimulated with a dynamic, binary white noise stimulus, and the spike-triggered average (STA) stimulus was computed for each RGC (Chichilnisky, 2001; Chichilnisky and Kalmar, 2002). The STA summarizes the spatial, temporal, and chromatic structure of the light response. The spatial RFs and time courses obtained from the STA were used to identify the distinct cell types, as described previously (Chichilnisky and Kalmar, 2002; Field et al., 2007; Rhoades et al., 2019).

### Electrical image (EI) computation

An electrical signature for each neuron on the array was obtained by computing the *electrical image* (or EI) (Litke et al., 2004). The EI of each cell (e.g. Fig. 1A) represents the average spatiotemporal pattern of voltage deflections produced on each electrode of the array during a spike from a given cell. A minority (∼3%) of spike clusters identified during spike sorting erroneously merged two different cells, and were excluded from analysis based on visual inspection. The remaining cells were used for subsequent analyses if their EIs featured spike amplitudes greater than twice the standard deviation of the recorded voltage trace (i.e. the electrical recording noise) at a minimum of three recording electrodes.

### Identifying cellular compartments and landmarks

As observed previously (Litke et al., 2004; Müller et al., 2015), the biophysical properties of somas, axons, and dendrites cause the spike waveforms recorded on MEA electrodes overlying each subcellular compartment to have distinct shapes (Fig. 1A, top row). Compartments were identified at each EI electrode in two steps. First, waveforms with a larger positive phase than negative phase were labeled as dendrites. Second, the waveforms at remaining electrodes were identified as somatic, mixed, or axonal, based on the ratio of the first positive peak to the second positive peak. Thresholds for each respective compartment were determined by two empirically observed inflection points (0.06 and 1.32 respectively) in the cumulative distribution of ratios-- across tens of thousands of spike waveforms obtained from thousands of recorded parasol and midget cells.

The identification of the cellular compartment at each electrode allowed for the estimation of two morphological landmarks for each RGC used in several analyses. The location of each RGC soma was estimated by taking the centroid of all identified somatic electrodes, weighted by the negative peak amplitude recorded on each electrode. The orientation of the axon was determined by computing a vector from the soma centroid to the nearest axonal electrode more than 180 μm away. The axonal-initial-segment (AIS) location was assumed to be 13 μm along this vector from the soma centroid (Sekirnjak et al., 2008). The axon midline was estimated by conducting amplitude-weighted fitting of a 3rd degree polynomial spline to all identified axonal electrodes.

### Electrical stimulation and spike sorting

Electrical stimuli consisted of triphasic current waveforms consisting of a cathodal phase flanked by two anodal charge-balancing pulses, with relative amplitudes 2:-3:1. Each phase was 50 μs in duration. The shape of the triphasic pulse was chosen to minimize the recorded electrical stimulation artifact (Hottowy et al., 2012). At each of the 512 or 219 electrodes on the 60 μm or 30 μm MEA respectively, triphasic pulses with thirty-nine stimulating-phase amplitudes increasing from 0.1 to 4.1 μA in 10% increments were delivered 25 to 50 times each. Spikes were elicited with sub-millisecond temporal precision (Jepson et al., 2013) by directly depolarizing the cell, as shown by synaptic transmission blockade (Sekirnjak et al., 2006). A semi-automated method was used to subtract electrical artifacts from the raw post-stimulation data and assign spikes to cells using waveform templates derived from their EIs, as described previously (Jepson et al., 2013; Mena et al., n.d.), supplemented by another, previously described semi-automated clustering spike-sorting method well-suited for analyzing electrodes with small peak waveforms (Madugula et al., 2020). To enable accurate spike assignment, at each electrode, only cells with recorded spike amplitudes of at least 25 μV were analyzed, because spikes with lower amplitudes were difficult to distinguish from background electrode noise. Stimulation amplitudes above axon bundle activation thresholds, which were estimated at each electrode using a previously described method for the 60 μm-pitch MEA (Tandon et al., 2021) or a custom algorithm for the 30 μm-pitch MEA (see below), were not analyzed in order to avoid potential off-array RGC activation. For each stimulus amplitude, the evoked spike probability was computed across repeats. The dependence of spike probability on stimulation current amplitude was modeled by a sigmoidal relationship:

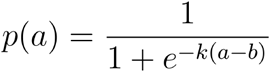

where a is the current amplitude, p(a) is the spike probability, and k and b are free parameters. Fitted sigmoidal curves were used to compute the *activation threshold* (b), defined as the current amplitude that elicits a spike with 50% probability, and *activation slope* (k). All fitted curves were manually inspected for goodness of fit, and a small minority (∼5%) were discarded due to either overfitting on spuriously identified spikes, or mistaken assignment of spikes to neighboring cells.

### Identification of axon bundle activation thresholds on the 30 μm MEA

Axon bundle activation thresholds on each electrode were determined by an automated method based on a previously described algorithm (Tandon et al., 2021), modified to avoid bias resulting from differences in array geometries, and from smaller axon spike amplitudes for central compared to peripheral RGCs. For each preparation, a threshold voltage was first determined to identify electrodes that recorded significant axonal signals in response to electrical stimulation, as follows. For each RGC recorded during white noise visual stimulation, the electrodes recording axonal signals were identified as described above and the average axonal spike amplitude was determined. The median axonal spike amplitude across all recorded RGCs was computed and was taken to be the threshold voltage. Next, to determine the axon bundle activation threshold, for each stimulus current applied, electrodes were first identified as either activated or inactivated, depending on whether the recorded signal was above the threshold voltage. Activity on the array was identified as an axon bundle activation event when the activated electrodes formed a contiguous path reaching at least two non-adjacent edges of the electrode array. The bundle activation threshold was then defined as the minimum current level at which an axon bundle activation event was evoked. Electrodes near the border of the array (outer two rings of electrodes) were excluded from analysis because their proximity to the edge complicates the ability to unambiguously distinguish RGC activity from axon bundle activity.

### Retinal preparation, cell, and activation curve selection

Of 53 total peripheral preparations from 50 different macaque retinas, 23 were included. Additionally, of 11 total central *raphe* preparations from 11 different macaque retinas, 10 were included. Central retina preparations were obtained from the *raphe* region, at 2-4.5 mm eccentricity along the temporal horizontal meridian, and recorded using an MEA with 30 μm electrode spacing (see above). Finally, of 3 total peripheral preparations from 3 different human retinas, 3 were included. In these preparations, the excised segment of retina covered more than 80% of the electrodes on the array, and the population RGC firing rate exhibited variance less than 20% of the mean. Only somatic and axonal cell-electrode pairs recording spikes greater than 25 μV, with activation curve thresholds less than both the bundle threshold on that electrode, which accounted for ∼80% of electrodes that were not able to be analyzed, and the upper current limit of the stimulation range (4 μV) were considered for analysis. Based on these criteria, 246 cell-electrode pairs from 59 ON and 43 OFF peripheral macaque parasol cells, 336 cell-electrode pairs from 185 ON and 70 OFF peripheral macaque midget cells, 189 cell-electrode pairs from 72 ON and 54 OFF *raphe* macaque parasol cells, and 16 cell-electrode pairs from 10 human parasol and 5 human midget cells were analyzed.

### Derivation and fitting of inference relations

The relationship between activation threshold and recorded spike amplitude, and the relationship between activation curve slope and threshold, were both modeled as inverse functions of the form 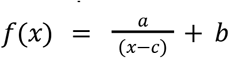. This functional form was chosen based on theoretical predictions from the literature (Rattay and Wenger, 2010), and the observation that normalizing sigmoidal activation curves by their thresholds resulted in curves with similar slopes within each cellular compartment. Inverse functions for each activation curve parameter were fitted by maximizing the log-likelihood of the evoked spike probability data. For simulated data (see below) however, the inverse function was fit directly.

### Biophysical simulations

RGC simulation was conducted using the NEURON software package (Finn et al., 2020; Hines and Carnevale, 2001) to create a model cell featuring five ion channel types, along with an extracellular stimulation mechanism, with an integration timestep of 0.001 ms. The properties of the model RGC were based on ion channel densities and compartment sizes adapted from voltage-clamp and morphology measurements of rat ON-alpha RGCs (Fohlmeister et al., 2010, 1990; Raghuram et al., 2019). Slight modifications to the model geometry were made to increase resemblance to primate peripheral parasol and midget RGCs, guided by previous measurements (Grosberg et al., 2017; Jeng et al., 2011; Kántor et al., 2018; Peterson and Dacey, 1998; Watanabe and Rodieck, 1989). The final parasol cell model consisted of a spherical soma with a diameter of 20 μm, a 40 μm long bent axon hillock with a diameter of 4 μm, a 40 μm long sodium-channel band (SOCB) axonal region with diameter tapered from 4 μm to 0.8 μm, a 90 μm long narrow axon compartment with diameter 0.8 μm, and a 2880 μm long distal axon compartment with a diameter of 1.2 μm (Fig. 5A). All compartments consisted of 5 μm finite-element segments. The final midget cell model consisted of a spherical soma with a diameter of 15 μm, a bent axon hillock with a diameter of 3.2 μm, a sodium-channel band (SOCB) axonal region with diameter tapered from 3.2 μm to 0.8 μm, a narrow axon compartment with diameter 0.8 μm, and a distal axon compartment with a diameter of 0.9 μm. The dendritic compartment morphology for both modeled cells was taken from (Kántor et al., 2018) (note the dendritic compartment is not shown in Fig. 5A due to its complex geometry).

The extracellular environment was assumed to be an isotropic, uniform medium with a linear resistivity of 1 kOhm-cm (Kasi et al. 2019; Hadjinicolaou et al. 2016). Extracellular stimulation was conducted with a collection of simulated point-electrodes located at a retinal depth of 7 μm relative to the axon, at 60 μm intervals along the length of the soma and axon (see Fig. 5A). The relative geometries between the electrodes and underlying modeled cell was varied to simulate various effective distances from the cell. Specifically, the electrode location was jittered along the axes parallel to the cell in steps of 5 μm, and along the axes perpendicular to the cell in steps of 2 μm. Simulated stimulus waveforms consisted of triphasic current pulses with shapes and amplitudes matching those used in *ex vivo* experiments (see above). Post-stimulation currents resulting from simulated action potential currents in the model cell were summed across each model element and used to compute simulated recording voltage at each stimulating electrode location.

The non-deterministic nature of extracellular stimulation putatively arises from stochastic membrane voltage fluctuations, caused by a combination of rapid ion channel state changes and synaptic inputs. To approximate these fluctuations, random voltage values drawn from a Gaussian distribution were added to the membrane voltage of model segments, at intervals of 0.15 ms. The noise was normalized to reflect the relative ion channel density differences between the axonal and somatic segments: noise with a standard deviation of 3 mV was injected at the somatic segments (F. Rieke, personal communication), versus noise with a standard deviation of 0.75 mV injected at the axonal segments. The resulting noise was smoothened by a 1D Gaussian across five neighboring segments at a time along the cell, in order to avoid unrealistically sharp changes in membrane voltage between neighboring model elements.

### Simulated image reconstruction

Simulation of perceived images resulting from application of stimulating current pulses, and selection of optimal electrical stimuli, were based on previous work (N. P. Shah et al., 2019) and consisted of several steps.

First, the contribution of RGC activity to visual perception was estimated in each of three retinal preparations, assuming that perception represents an optimal linear reconstruction of the stimulus from the RGC responses (Brackbill et al., 2020; Warland et al., 1997). Optimal linear reconstruction was examined by simulating the response of each cell to 10,000 12×6 random black and white noise training images, based on the spatiotemporal STA (see above), and then computing *reconstruction filters* from the simulated responses for each cell as follows. The simulated nonnegative response to each training image was estimated by taking the inner product of the STA and that image, and rectifying the result. Then, the reconstruction filter was obtained by performing least squares regression of the simulated responses against the set of training stimuli.

Next, for each of 15 randomly generated 12×6 black and white images, 40 electrical stimuli were chosen to (approximately) produce an appropriate neural response in each retinal preparation. Each stimulus consisted of a particular current amplitude on a particular electrode and produced spikes with probabilities from 0 to 1 in one or more RGCs, estimated using either measured, inferred, or fixed activation curves. The fixed curve was assumed using parameters of 1.2 μA for threshold and 12.1 μA^-1^ for slope, obtained by averaging across all measured responses to electrical stimulation in 53 preparations (see Fig. 6 B-G). For each type of activation curve, stimuli were chosen so that the sum of their presumed contributions to perception, based on the linear reconstruction filters weighted by the expected response, minimized the normalized mean squared error (NMSE) between the stimulus and reconstructed image. Normalization was relative to the highest achievable mean-squared error across images and retinas, for ease of interpretation. The number of electrical stimuli (40) was empirically determined as the value beyond which, on average, reconstruction using inferred activation curves no longer reduced the NMSE.

To quantify the effectiveness of using inferred over fixed activation curves for image reconstruction, NMSE values of reconstructed images based on the three curve types were collected for each retinal preparation and target image. Then, NMSEs for inferred and measured activation curves were compared to NMSEs for fixed curves (see Fig. 6B). Across preparations and targets, systematic deviations from the x=y line, which corresponds to using the fixed curve, were captured by slopes obtained from linear regression. Finally, the overall value of performing reconstruction with inferred activation curves was given by |(s_m_ - s_i_)/(s_m_ - 1)|, where s_m_ is the slope obtained with measured activation curves and s_i_ is the slope obtained with inference.

